# Comparative miRNA Transcriptomics of Mouse and Macaque Reveals MYOC is An Inhibitor for *C. neoformans* Invasion into Brain

**DOI:** 10.1101/2022.02.14.480319

**Authors:** Hailong Li, Xiaoxu Han, Wei Du, Yang Meng, Yanjian Li, Tianshu Sun, Qiaojing Liang, Chao Li, Chenhao Suo, Xindi Gao, Yu Qiu, Wen Tian, Minghui An, Hui Zhang, Yajing Fu, Xiaolin Li, Tian Lan, Sheng Yang, Zining Zhang, Wenqing Geng, Chen Ding, Hong Shang

**Author notes:** **Correspondence:** Hong Shang, Chen Ding.

## Abstract

Cryptococcal meningoencephalitis is an emerging infection shifted from primarily ART-naive to being ART-experienced HIV/AIDS patients, COVID-19 patients and also in immune competent individuals, mainly caused by the human opportunistic pathogen *Cryptococcus neoformans*, yet mechanisms of the brain or CNS dissemination remain to elucidate, which is the deadest process for the disease. Meanwhile, illustrations of clinically relevant responses in cryptococcosis were limited, as the low availabilities of clinical samples. In this study, macaque and mouse infection models were employed and miRNA-mRNA transcriptomes were performed and combined, which revealed cytoskeleton, a major feather in HIV/AIDS patients, was a centric pathway regulated in both two infection models. Notably, assays of clinical immune cells confirmed an enhanced “Trojan Horse” in HIV/AIDS patients, which can be shut down by cytoskeleton inhibitors. Furthermore, we identified a novel enhancer for macrophage “Trojan Horse”, myocilin, and an enhanced fungal burden was achieved in brains of MYOC transgenic mice. Taking together, this study reveals fundamental roles of cytoskeleton and MYOC in blocking fungal CNS dissemination, which not only helps to understand the high prevalence of cryptococcal meningitis in HIV/AIDS, but also facilitates the development of novel drugs for therapies of meningoencephalitis caused by *C. neoformans* and other pathogenic microorganisms.

## Introduction

Fungal invasive diseases are an increasing threat to global public health, lead to more than one million deaths every year, mainly caused by species of *Candida albicans, Aspergillus fumigates* and *Cryptococcus neoformans*. Cryptococcal meningoencephalitis (CM) was the leading diseases among fungal meningoencephalitis, caused 181,000 deaths annually, with mortality rates of 100% if untreated[1]. Recent studies showed that the prevalence and mortality has not been decreased, and the appeals for actions on CM was raised[2,3]. Additionally, CM is a major risk for HIV/AIDS patients with 77%-90% prevalence, leading to about 15% deaths of HIV/AIDS patients annually[4]. In the era of ART (antiretroviral therapy), CM is also an emerging infectious disease, shifted from primarily ART-naive to more than 50% being ART-treated[2]. Seriously, morbidity of cryptococcosis in immune competent is increasing rapidly in China, Australia, Canada, and other countries and regions[4–8]. Importantly, COVID-19 patients suffered from secondary cryptococcus infections, suggesting cryptococcosis is an important issue in the post-COVID-19 era [9–12]. Fungal central nervous system (CNS) dissemination is the lethal procedure, which is vital for both fungal colonization and fungal clearance, but lack of mechanisms from in vivo or clinical studies.

To date, massive researches have focused on CNS invasion during fungal meningitis and numerous milestones achieved, including the facts that cryptococci interact with brain epithelial cells directly and indirectly by microscopy or electron microscope[13–16]. These researches are mainly performed on cell lines, mice, zebra fish or rabbits[17], However, clinically relevant evidences were limited, as the unavailability of clinical samples. Cryptococcal meningoencephalitis are predominant prevalent among immune compromised patients, especially HIV/AIDS patients [18–20]. Mechanisms from clinical HIV/AIDS patients were the most direct approach to cryptococcus meningoencephalitis. Dysfunctional cytoskeleton of immune cell was one of pathological features in HIV/AIDS patients[21]. Meanwhile, previous studies indicated that *C. neoformans* infections also disturbed cytoskeleton pathways in mice and human brain microvascular endothelial cells[22,23]. No evidence has revealed the mechanisms between dampened cytoskeleton and cryptococci brain dissemination. Identification relationships between the cytoskeleton and fungal infections may provide novel mechanisms of fungal pathogenesis and targets for drug development in mycosis.

Furthermore, responses of hosts during infectious diseases are often divided into DNA, mRNA and protein levels, and also, the processes of post transcriptional and translational modifications[22,24,25]. Previous studies identified many responses at mRNA, protein and post translational levels, including key pathways and modulators, such as mineral metabolism, IL-17 signaling pathway, sugar metabolisms, OCSTAMP, IL-5, IL-13 and IL-17A [22,24,26–31]. Post transcriptional modifications, assumed by miRNAs, also play important roles in mRNAs and proteins biosynthesis. Recently, several studies have demonstrated key functions of non-coding RNAs during *C. neoformans* infections in vitro, However, little was known in vivo[32–35].

In this study, to mimic human responses, macaques were employed, and in vivo miRNA-mRNA network was constructed. We found that cytoskeleton pathway is the core pathway regulated by *C. neoformans* in both macaque and mice. Moreover, assays of clinical immune cells confirmed an enhanced “Trojan Horse” in HIV/AIDS patients, and intervention of cytoskeleton pathway was able to disturb “Trojan Horse”. Furthermore, cytoskeleton associated gene, *MYOC*, were identified as important factors for “Trojan Horse” by THP-1 cells and transgenic mice. Collectively, these findings demonstrate global responses at miRNA-mRNA regulatory level, reveal novel modulators for fungal invasion and benefit novel therapies for fungal infectious diseases.

## Materials and Methods

### Ethics statement

All work with human cells was reviewed and approved by the Medical Research Ethics Committee of the First Affiliated Hospital of China Medical University (2021-63-2). Animal infection experiments in macaques and mice were reviewed and ethically approved by the Research Ethics Committees of the College of Life and Health Sciences of Northeastern University (16099M) and Wincon TheraCells Biotechnologies Co., Ltd. (WD-20150701-a). All animal experiments were carried out according to the Guide for the Care and Use of Laboratory Animals issued by the Ministry of Science and Technology of the People’s Republic of China.

### Animal Infection

Macaques and mice were purchased from Grandforest Co., Guangxi, China and Changsheng Biotech, China, respectively, and infections were performed same to the previous article[22]. Briefly, six female macaques were divided into two groups. Monkeys were anesthetized by ketamine (10 mg/kg) intraperitoneally injection (IP), and then infected via intratracheal injection with 10^8^ cells/kg *C. neoformans* H99 cells. Controls were infected using the same volume of PBS. Mice were anesthetized and infected intranasally with 10^5^ fungal cells. Macaque and mice were monitored for signs of infection and humanely killed at day 7 or 14 post infections, or used for survival rates detections.

### miRNA sequencing and analysis

Total RNA was isolated using TRIzol. Assessments of RNA were done using a NanoDrop 8000 spectrophotometer. Small RNA-seq libraries were prepared by using TruSeq^®^ Small RNA Library Prep Kit according to the manufacturer’s protocol. miRNA libraries construction and sequencing were entrusted to Shanghai Personal Biotechnology Co., Ltd. (China), and then single-end sequencing was conducted by Illumina NextSeq 500 platform. Raw data were obtained, clean and unique reads were mapped to corresponding genomes by bowtie. Expression of miRNAs were identified by using quantifier.pl in Mirdeep2 based on miRBase21. Differentially expressed miRNAs were calculated by DESeq.

### Quantitative PCR of miRNAs

Total miRNAs were isolated by miRNeasy Mini Kit (QIAGEN) according to the manufacturer’s manual. First strand cDNA was synthesized using random oligonucleotides and miRcuRY LNA Univerersal RT microRNA PCR Universal cDNA Synthesis Kit ? (EXIQON, USA), U6 was employed as an internal reference. Primers (Table S4) were designed by using miRprimer and the best primer pairs were selected[36]. RT-qPCR was performed by miRcuRY LNA Univerersal RT microRNA PCR Exilent SYBR master mix (EXIQON, USA)) and StepOne Plus.

### Histopathology, colony forming units (CFU) and survival rates assessments

Lung tissues were collected from macaque and mice, as described previously [22]. For histopathology analyses, tissue samples were fixed with paraformaldehyde, frozen, and processed using a cryostat microtome (CM1850; Leica). Tissue sections of 10 μm in thickness were stained with mucicarmine or hematoxylin/eosin. For CFUs determination, homogenized lung and brain tissues were diluted and plated onto YPD plates and then incubated at 30°C for two days. Colonies were counted and calculated. For survival rates detection, C57BL/6 mice were divided randomly. Body weight was examined. Fifteen percentage decrease of initial body weight was identified physiological endpoint.

### Construction of MYOC overexpressed THP-1 cell line and THP-1 derived macrophages (TDMs) differentiation

MYOC gene was cloned into pCDH-EF1*α*-MCS-T2A-Puro plasmid. Lentivirus was packaged by using jetPRIME^®^ DNA & siRNA Transfection Reagent (polyplus) according to instructions. Infections of THP-1 were performed by centrifugation at 1200 × g at 37 °C in 24-well plates for 2h. Screening of puromycin resistance was conducted 72h post transduction for 2 weeks at 5 μg·mL^-1^. TDMs were differentiated for 48h in RPMI-1640 medium (10% FBS) containing 1% penicillin/streptomycin and 250 ng/mL PMA (Sangon Biotech, China).

### Human monocytes isolation and monocyte derived macrophages (MDMs) induction

PBMC were isolated by Ficoll-Paque PLUS density gradient media (Cytiva). Blood samples were centrifuged in a swing bucket rotor at 400 × g for 30 minutes at 25 °C with acceleration set at 5 and break at zero, and followed by purification in cold PBS for 2 times and centrifuged at 300 × g for 10 minutes at 4 °C with acceleration and break set at 5. 1×10^6^ PBMC were seeded onto 48-well plates in RPMI-1640 media without FBS for 1h to stick monocytes, and fresh RPMI-1640 medium (10% FBS) containing 1% penicillin/streptomycin and 50 ng/mL M-CSF was exchanged, and fresh medium was renewed at days 3 and 6 during monocytes differentiation. MDMs were ready to use on day 7.

### Phagocytosis effectivity, killing and transmigration assessments

MDMs or TDMs were seeded on 48-well plates at 100,000 cells per well 24h before fungi interaction. GFP-expressed strains (H99) were incubated overnight at 30 °C, washed, opsonized by 18B7(1mg·L^-1^) at room temperature for 30 min. Incubate fungal cells with macrophages in CO2 incubator overnight at MOI 1:10. Macrophages and fungal cells were washed 5 times with PBS buffer thoroughly and then digested by trypsin. Phagocytosis effectivity was detected by flow cytometry. For killing tests, after fungi and macrophage incubation (24h), supernatant was collected and cells were washed and collected. Total lysates and supernatant were diluted and plated on YPD agar plates and colonies were counted and calculated 48h post incubation at 30°C. Migration of THP-1 and MDMs were performed by trans-well.

### Construction of MYOC transgenic mice

MYOC transgenic mice were produced by Beijing View Solid Biotechnology, China. The linear plasmid pCAG-MYOC cut by the *BstEII* restriction enzyme (NEB) was purified, which were injected into zygotes of C57BL/6 mice in M2 media (Millipore) using a FemtoJet micromanipulator (Eppendorf, Germany). Microinjected zygotes were transferred into pseudo pregnant female mice. All mice were maintained in a specific pathogen-free facility. Genotype identification was performed by PCR and sequencing from 2-week-old newborn mice with primers (Table S4). Transgenic mice were mated with wild-type C57BL/6 mice to obtain heterozygous mice and colony expansion.

### Bioinformatics Analysis

Targeted genes of miRNAs were predicted using miRWalk 3.0 [37]. Regulatory network miRNA-mRNA was constructed by Cytoscape. Gene ontology and KEGG analyses were performed using R version 4.1.2, clusterProfiler v4.2.0, org.Hs.eg.db version 3.14.0, org.Mm.eg.db version 3.14.0 packages, and plotted by ggplot2 version 3.3.5. KEGG of miRNAs was performed by using a web-based application named miEAA [38] and plotted by ggplot2. Heatmap of expression was generated by Pheatmap version 1.0.12. Homology analysis was performed based on miRbase search engine database (https://www.mirbase.org/search.shtml).

### Statistics and reproducibility

All experiments were performed at least biologically triplicated to ensure reproducibility. Statistics of phagocytosis effectivity, RT-qPCR, CFU were calculated using GraphPad Prism 9.0. An unpaired or paired student *t* test was performed. When the *p*-value was less than 0.05, statistical significance was recognized.

## Data and Software Availability

The RNA-seq raw data files have been deposited in NCBI’s Gene Expression Omnibus (GEO) with GEO Series accession ID GSE122785 previously[22]. Raw data of microRNA-seq was ready and open to researchers and can be provided upon request.

## Author Contributions

H.S. and C.D. conceived the project. H-L.L., X-X.H., W-Q.G. and C.D. designed the study. T.L., S.Y., and C.D. performed the monkey infection experiments. H-L.L., W.D., Y.M. Q-J.L. Y-J.L. and T-S.S. performed mouse infection experiments. H-L.L., X-L.L., M-H.A., Y.Q., H.Z., X-D.G. and Z-N.Z. participated in data analysis. C-H.S. and C.L. performed histopathology staining. H-L.L., W.T. and Y-J.F. performed cell experiments. H-L.L. X-X.H., W-Q.G., C.D., and H.S. composed the manuscript.

## Results

### MicroRNA transcriptomics in macaque and mouse during cryptococcal pneumonia

We have previously unveiled responses in macaque and mouse at transcriptional level and identified several regulators and pathways during *C. neoformans* infections[22]. To investigate global responses at post transcriptional level, microRNA transcriptomes were performed using lung tissues isolated from macaque and mouse infection models (Figure 1), and miRNA-mRNA network was constructed, which gave a comprehensive response in cryptococcosis clinically relevant.

**Figure 1.**
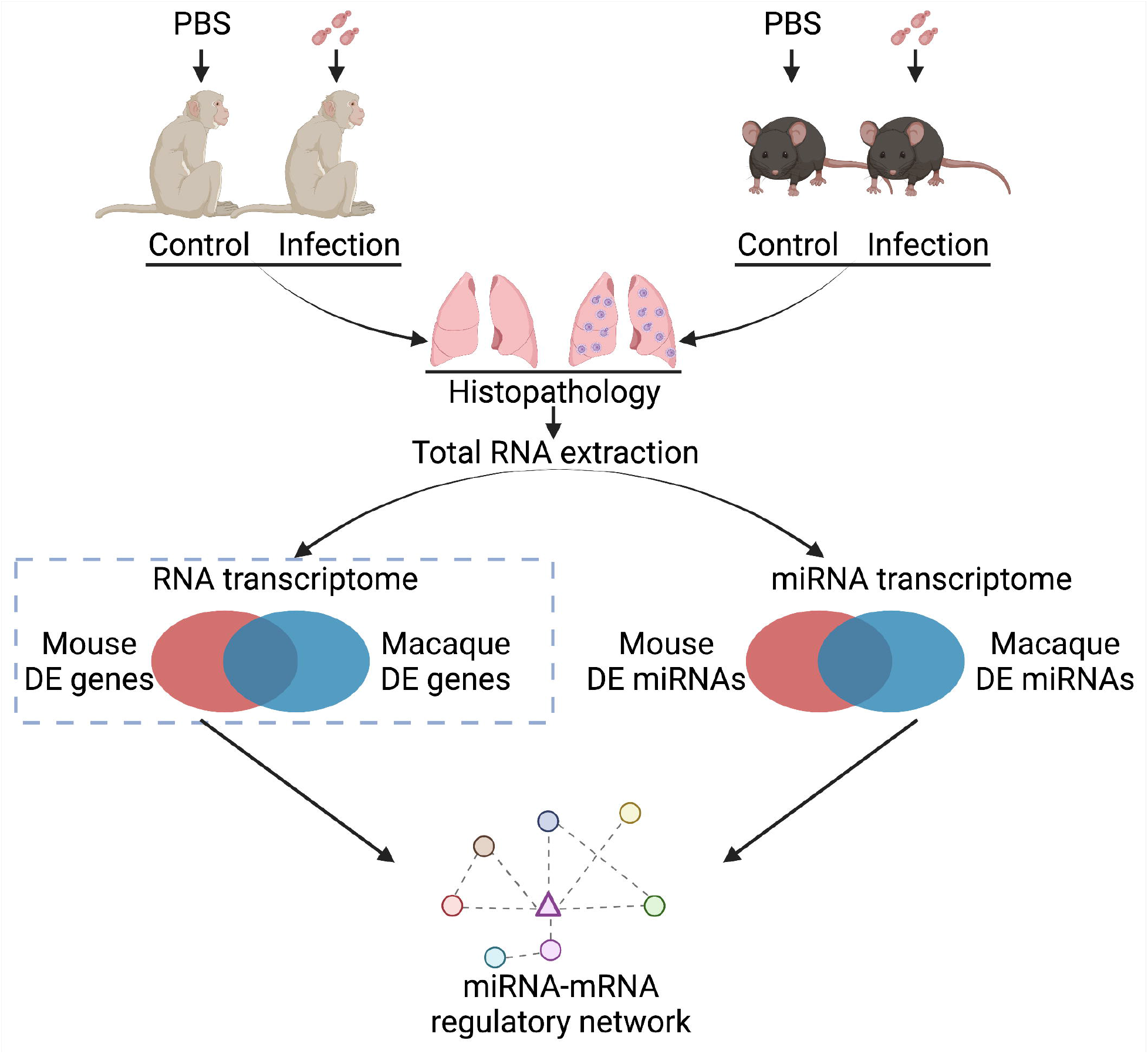
Flow chart for animal infections and RNA sequencing. Six macaques and mice were divided into two groups randomly and infected by H99 intranasally. Lung tissues were isolated 7 days post infections. Total RNAs were isolated for mRNA-seq and miRNA-seq followed by confirmation of histopathology. Differentially expressed miRNA-mRNA regulatory network was constructed. Figure 1 was created with BioRender.com

Pathologies of macaque and mouse lung tissues were confirmed by histopathology observations using mucicarmine and hematoxylin/eosin staining, showing the capsular structure *of C. neoformans* (Figure2 A, B). Total RNAs were extracted and miRNA transcriptomes were performed. Total and unique reads from omics were calculated, with a number of more than 10^7^ of total read and 10^6^ of unique read (Figure S1A). Read number around 22 nucleotides were the most abundant (Figure S1 B, C). The clean read files were used to map the corresponding genome based on species specificity in miRBase21. However, macaque has few miRNA annotations, as which miRNA annotations of humans were employed for macaque. As a result, there are 1038 and 1166 expressed miRNAs in lung tissues from macaques and mice (Table S1). Principal component analyses (PCA) were analyzed by using read number of each detected miRNA, as shown in Figure 2 C, D, both infected and uninfected samples were reproducible. Differentially expressed miRNAs were performed by DESeq. Heatmaps of differentially expressed items were generated, with 32 miRNAs (4 down regulation and 28 up regulation) in macaque and 29 (5 down regulation and 24 up regulation) in mouse, shown as two clusters by column (Figure 2 E, F).

**Figure 2.**
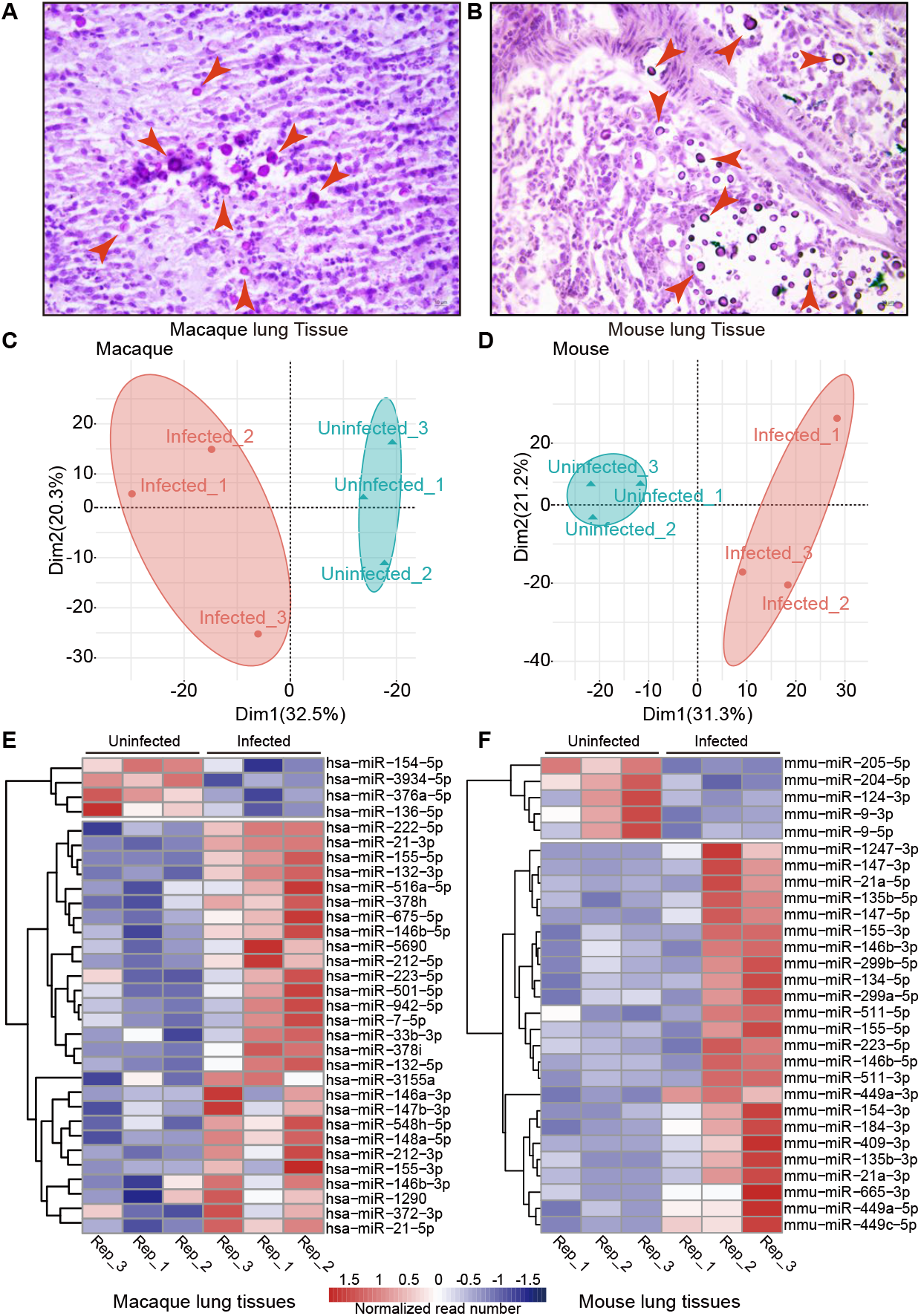
miRNA-Seq of mice and macaques in response to *C. neoformans*. **A B. Histopathology observation of infected macaque and mouse lung tissues**. Lung tissues from macaque and mouse were fixed and sectioned at 10 μm thickness and stained by using mucicarmine. Red arrows indicate *C. neoformans* cells, scale bar = 10 μm. **C D. PCAs of miRNA-Seq data**. **E F. Heatmaps of differentially expressed miRNAs**. miRNAs with *p*-value ≤ 0.05, fold change≥ 2 were considered as differentially expressed.

### Core responses of host derived from miRNA-mRNA Integrative analyses

To mine and mimic responses in human cryptococcosis extremely, homology analyses were performed (Figure 3A). Eight miRNAs were co-regulated in macaque and mouse during *C. neoformans* infections. Information about the 8 miRNAs were displayed, including IDs, foldchanges and homologous e-values, and RT-qPCRs were performed in mouse for the 8 miRNAs, which were consistent with the miRNA-Seq data (Figure 3 B, C, D).

**Figure 3.**
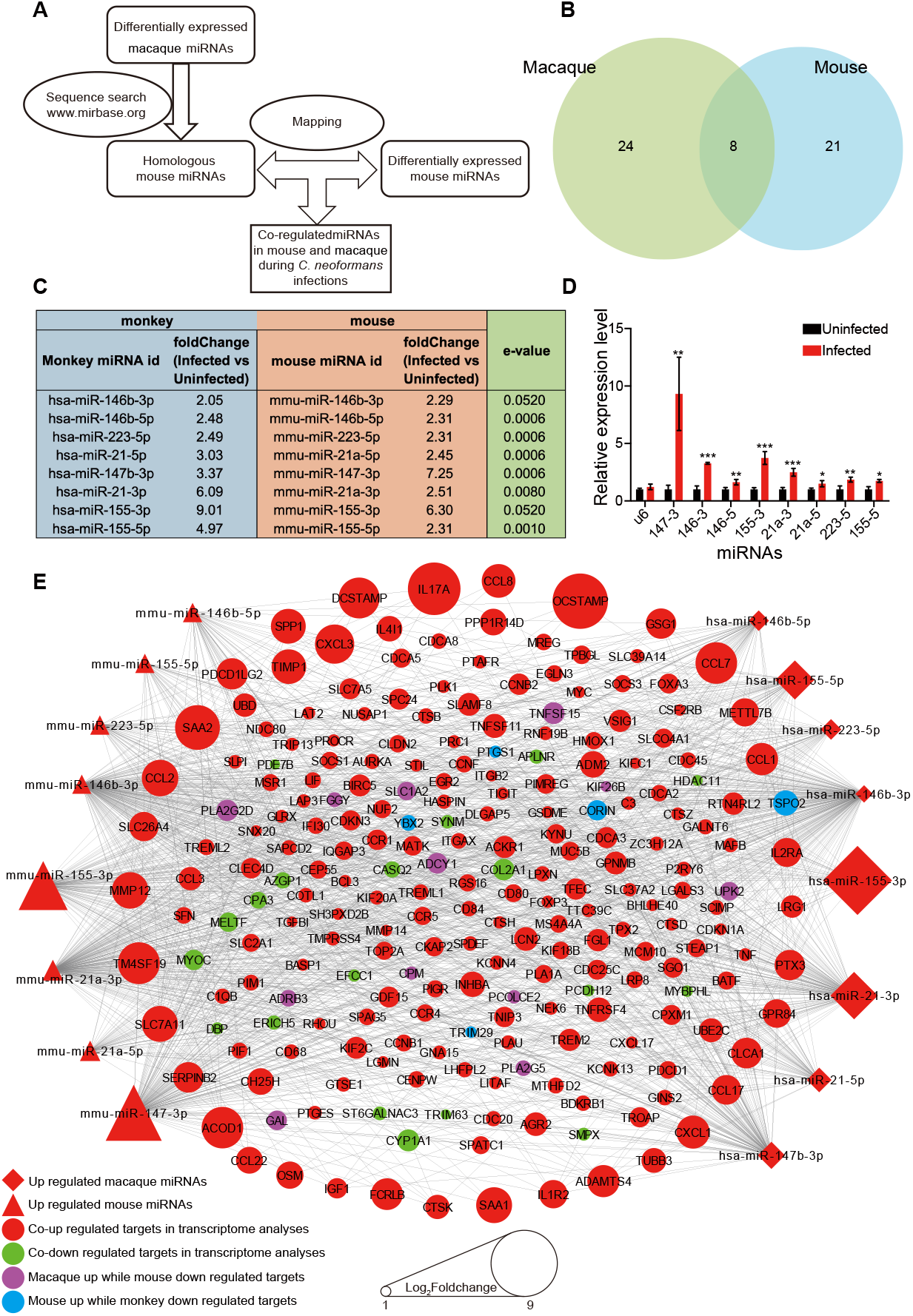
Core regulatory machinery of miRNA-mRNA network during *C. neoformans* infections. **A. Schedule of identification of homologous miRNA**. miRNAs of macaque were used for searching homologous miRNAs in miRbase. **B. Venn diagram of differentially expressed miRNAs**. **C. Information of 8 co-regulated miRNAs**. **D. RT-qPCR of the 8 co-regulated miRNAs in mouse**. Three biological replicates were performed. Mean and SEM were shown. Unpaired student *t* test was performed. \**p* < 0.05, ** *p* < 0.01, or \*\*\**p* < 0.005. **E. miRNA-mRNA core regulatory network**. Differentially expressed target genes of 8 co-regulated miRNAs were shown. The whole network is shown in Figure S3.

Targets of the 8 co-regulated miRNAs were predicted, then mapped to RNA-Seq, and omics of miRNA and mRNA were integrated (Figure S2). The core regulatory target mRNAs, co-regulated by the 8 co-regulated miRNAs, were selected for a mini miRNA-mRNA network, with 233 target mRNAs identified, including OCSTAMP, DC-STAMP, IL17A TNF and LIF, which demonstrated anti-microbial activities (Figure 3E). KEGG and GO analyses were performed of the 8 co-regulated and 223 co-regulated targets (Figure 4, Figure S3, Table S2 and Table S3). KEGG pathways associated immune system, lung diseases, and infectious diseases were significantly enriched in both miRNA and mRNA level, such as TNF signaling pathway, Cytokine-cytokine receptor interaction, Osteoclast differentiation, small cell lung cancer, non-small cell lung cancer and Influenza A (Figure S3). GO analyses identified terms involved cytokine activity, histone, immune cells, monocyte, myeloid leukocyte and cell cycle were significantly varied by *C. neoformans* (Figure 4). Interestingly, actin binding, microtubule and their associated complex, which constitute cytoskeleton system, were highly enriched in GO analyses, including cellular component, molecular function and biological process. These results indicated a potent function of cytoskeleton in cryptococcosis.

**Figure 4.**
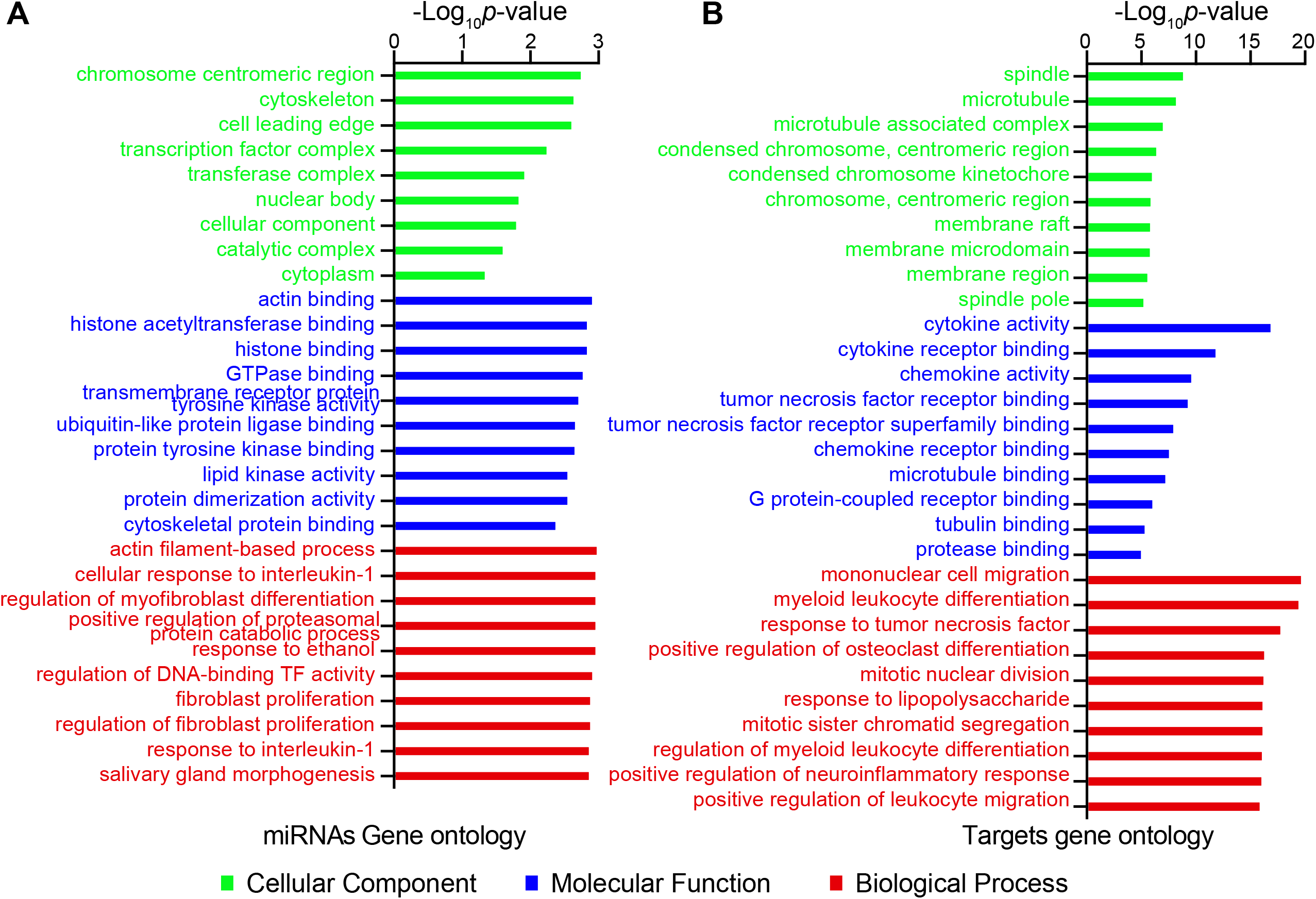
GO analyses of co-regulated miRNAs and mRNAs in macaque and mouse during *C. neoformans* infections. **A. GOs of miRNAs calculated by miEAA. B. GOs of target mRNAs enriched by Clusterprofiler**. Eight co-regulated miRNAs and co-regulated mRNAs from Figure 4E were employed for GO analyses, respectively. Top 10 or all significantly enriched GO terms were plotted. Green, blue and red columns represent cellular component, molecular function and biological process, respectively.

### “Trojan Horse” was enhanced in HIV/AIDS patients and can be dampened by cytoskeleton pathway inhibitor

Dysfunctional cytoskeleton of immune cell was one of pathological features in HIV/AIDS patients, who are the predominately population for cryptococcal meningoencephalitis. Based on PSM (Propensity Score Matching, PSM), 100 HIV patients and 200 healthy individuals were compared, number and percentage of monocytes were significantly enlarged in HIV patients (Figure 5B). To reveal the functions of host cytoskeleton during the battle between host and *C. neoformans*, phagocytosis and transmigration of MDMs from 9 HIV/AIDS patients and 8 healthy volunteers were compared (Figure 5A). As shown in Figure 5C, effectivities of phagocytosis were enhanced in HIV/AIDS patients, and more interestingly, the lifted MFI indicated more fungal cells were phagocyted or more intracellular proliferation in HIV/AIDS. Transmigration of MDMs has no change between HIV and healthy volunteers (data not shown), but positively correlated to capacities of phagocytosis (Figure 5D).

**Figure 5.**
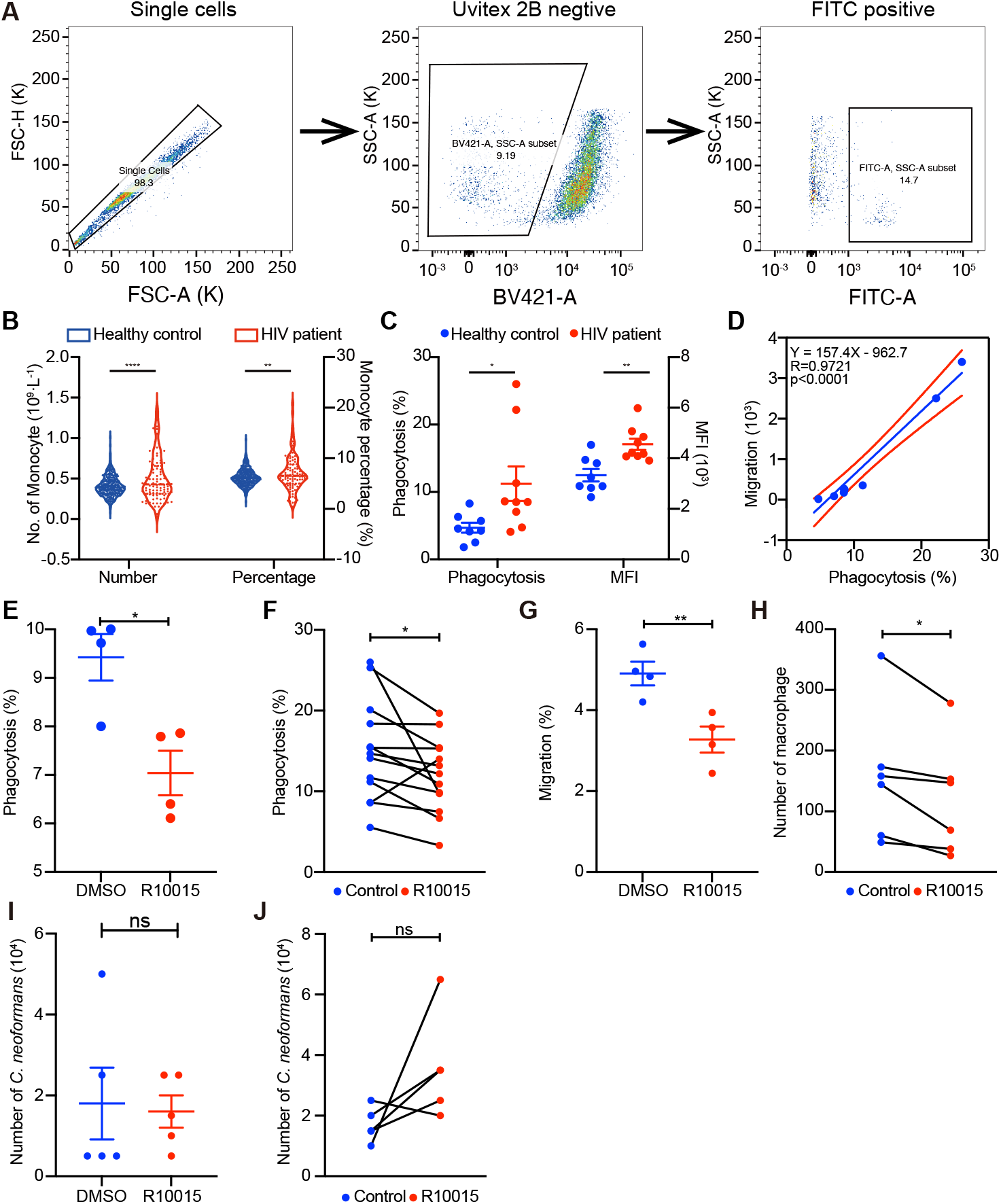
“Trojan Horse” was enhanced in HIV/AIDS patients. **A. Strategies for phagocytosis assessment by flow cytometry**. Cryptococcal internalization was determined by flow cytometry using GFP- expressed H99. During flow cytometry, single cells were selected, cells with negative Uvitex 2B were considered as phagocytes, and in which FITC positive cells were fungi internalized (Uvitex 2B -/ FITC +). **B**. Number and percentage of monocytes between HIV patients and healthy individuals from clinical data **C**. Phagocytosis effectivity of MDMs from HIV patients and healthy individuals **D**. Correlation of phagocytosis and migration in MDMs from HIV patients. **E, F**. Phagocytosis effectivity was inhibited by R10015 in TDMs(E) and MDMs(F) **G, H**. cells migration was dampened by R10015 in THP-1(G) and MDMs(H). **I, J**. R10015 does not affect killing *C. neoformans* in THP-1(I) and MDMs(J).

In order to confirm the association of phenomenon from HIV/AIDS patients with cytoskeleton, R10015, a cytoskeleton pathway inhibitor, was employed, which targets LIM Kinase (LIMK) powerfully in cytoskeleton pathway[39]. Based on the growth curve, 14.815 μM was selected (Figure S4A). THP-1 derived macrophages and human monocyte derived macrophages were employed and pre-treated by R10015 for 2h, washed and then incubated with opsonized *C. neoformans* overnight. As shown in figures, R10015 inhibited capacities of phagocytosis and migration of macrophages derived from both THP-1 cell lines and primary monocytes from human (Figure 5 E, F and G, H), however, R10015 did not affect the killing in TDMs in and MDMs (Figure 5 I, J). These results confirmed the vital roles of cytoskeleton in macrophage “Trojan Horse”.

### MYOC is an inhibitor for cryptococci brain dissemination by modulating macrophage “Trojan Horse”

To identify functions of cytoskeleton during *C. neoformans* infections, genes associated with cytoskeleton were screened, and myocilin was selected, a tubulin binding protein, encoded by MYOC gene, which was one of the centric modulators in miRNA-mRNA regulatory network and down regulated significantly during *C. neoformans* infections in both macaque and mouse (Figure 6 A, B). Meanwhile, MYOC was elevated in HIV/AIDS patients (Figure 6C). Previous studies revealed the protein was involved with cell migration and adhesion[40].

**Figure 6.**
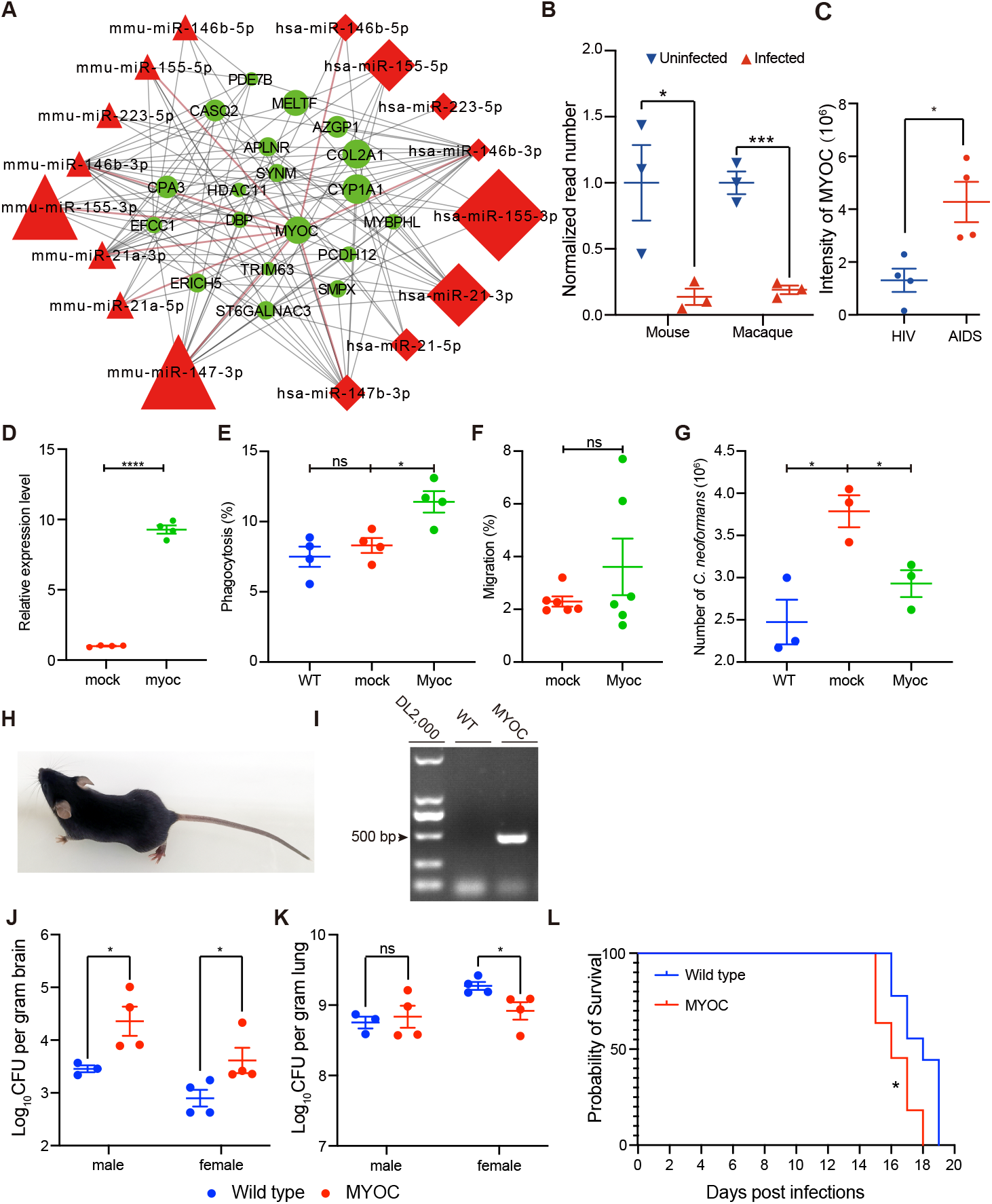
MYOC enhanced *C. neoformans* brain invasion by modulating macrophage “Trojan Horse”. **A**. miRNAs-mRNA network during *C. neoformans* infections. **B**. Normalized MYOC reads in mouse and macaque in response to *C. neoformans*. **C**. Intensity of MYOC protein from comparative proteomes between HIV and AIDS patients. **D**. Relative expression level of MYOC in MYOC gene edited THP-1 cell lines. **E. Effectivity of phagocytosis in MYOC overexpressed THP-1 derived macrophages by flowcytometry**. Four biological replicates were performed. Unpaired Student *t* test was performed. * *p* < 0.05. **F**. Capacity of migration in MYOC overexpressed THP-1 cell lines. **G**. Killing tests of MYOC overexpressed THP-1 derived macrophages. **H. Photo of MYOC transgenic mice**. The cute picture is one of our MYOC transgenic mice,5-week-old, which showed healthy and eye-functioned. **I. MYOC gene fragment was checked by agarose electrophoresis in MYOC transgenic mice**. Genome DNA was isolated and PCR and electrophoresis were performed. Target PCR products was 498 bp. **J, K. CFU assessments of MYOC transgenic mice, brain(J) and lung(K) tissues**. Seven MYOC transgenic mice (3 male and 4 female) and 8 wild type mice (4 male and 4 female) from the same cages were used for CFU assays. Unpaired Student *t* test was performed. * *p* < 0.05. **L. Survival tests of MYOC transgenic mice**. Eleven MYOC transgenic mice (5 male and 6 female) and 9 wild type mice (3 male and 6 female) from the same cages were used for survival assays. *Log-rank* (Mantel-Cox) test was employed for statistical analysis. * *p* < 0.05.

To explore functions of MYOC, MYOC overexpressed THP-1 cell lines were constructed and mRNA levels of MYOC were quantified by RT-qPCR, which showed a 10-fold induction compared to mock cells (Figure 6D). Phagocytosis, migration and killing of macrophage cells in MYOC gene-edited cell lines were examined (Figure 6 C, E, F, G). Phagocytosis effectivity was increased significantly in TDMs when MYOC was overexpressed (Figure 6E). Killing capacity was also enhanced by MYOC overexpression, as shown in Figure 6G, less lived *C. neoformans* were detected by CFU assays. However, no changes were observed in migration assays (Figure 6F).

To evaluate functions of MYOC in vivo, MYOC-transgenic mouse was generated and employed (Figure 6H), which was confirmed by PCR (Figure 6I). Six-eight weeks old mice were used for CFU assessments and survival rates. To our surprise, CFU of brains was enhanced in MYOC transgenic mice, either in male or female groups (Figure 6J), however, lung CFU was decreased in females, while no changes in the male group (Figure 6K). Consistent with CFU assays, MYOC transgenic mice showed reduced survival times compared to wild type mice observed in survival rate tests (Figure 6L, S4B).

## Discussion

Cryptococcal meningoencephalitis is an emerging disease with high mortality, even under the ART conditions currently, and deserved more attention in post-COVID-19 era [1–3,9–12]. Brain dissemination is the lethal procedure. However clinically relevant mechanisms were limited[13–17]. In this study, we demonstrated the landscape at transcriptional and post transcriptional levels in mouse and macaque infection models, which were employed for mimicking and unveiling responses at miRNA-mRNA regulatory levels in humans. Previously studies indicated mouse was the most used animal model, however, varied from humans a lot, such as the process of immune cells maturation. Our data from *M. fascicularis*, 92.83% sequence identity to human, a little more clinically relevant, may serve as a database for cryptococcosis or mycosis. To investigate core responses during *C. neoformans* infections, miRNA-mRNA combined, GO and KEGG analyses were performed. Eight key miRNAs were identified in our omics, such as miR-146a, miR-223 and miR-155, which were induced in monocytes by co-culture with *C. neoformans* in vitro[32]. We did not focus on the functions of unique miRNAs, and we characterized cytoskeleton pathway as a core regulatory modulator based on enrichment analyses of miRNAs and their targets. Previous studies identified *C. neoformans* disturbed cytoskeleton when interacted with brain endothelial cells[23].

Our data identified the fundamental roles of cytoskeleton during “Trojan Horse” formation, and provided potential targets for novel “orphan drug” development. Studies have indicated the relationship between cytoskeleton and fungal infections[23,41–44]. However, no mechanisms and in vivo studies to conform. Previous studies highlighted important roles of “Trojan horse” by macrophages during fungal CNS invasion, and the processes are cytoskeleton dependent. Studies have demonstrated migration, adhesion and phagocytosis are key features for phagocytes[45]. Our studies illustrated the basic functions of cytoskeleton dynamics on capacities of phagocytosis and migration of macrophages, which can be shut down by cytoskeleton inhibitors, However, small molecule drugs, such as vanadate, cytochalasin D, Y27632 and R10015 are mainly toxic to human [23,39,46]. The toxicity involves fundamental roles of innate immunity and acquired immunity of macrophages, NK cell and T cells, who also play important roles during fungal, bacterial and virus infections. Overcoming toxicity of cytoskeletal inhibitors is the main barrier to clinical application.

Further analyses revealed that dysfunctional cytoskeleton may account for the high prevalence of cryptococcal meningitis in HIV/AIDS patients. Studies indicated a similar variation of cytoskeleton pathway during HIV infection and *C. neoformans* or *A. fumigates* infections [23,42,43,47,48]. Indeed, *C. neoformans* is an environmental yeast globally ubiquitous, and easily accessible for all individuals[49]. However, why do these happen much more in HIV/AIDS? Previous studies suggested the decreased number of CD4 T cell was a high risk, however contradictory to the high prevalence of high level CD4 T cells who are ART experienced[50–52]. Another hypothesis is the high exposure to *C. neoformans* environment, but, inconsistent with the low prevalence of other people in the same environment[53]. Our data suggest the dampened cytoskeleton structure maybe the reason. In HIV/AIDS patients, numerous studies have proved cytoskeleton was dampened in immune cells of PBMC[39,48,54], which contributes to fungal cells CNS invasion by “Trojan Horse”. Furthermore, a recent study proved vomocytosis was enhanced by HIV infection in macrophage[55], which process was also regulated by the cytoskeleton.

We identified MYOC as a novel modulator for fungal invasion by regulation on macrophages. Our data showed a repression of MYOC in *C. neoformans* infections and a reduction in HIV infections. Myocilin was proved co-localized with microtubules, endoplasmic reticulum (ER), and Golgi apparatus[56,57] and overexpression of MYOC induces a loss of actin stress fibers[58]. Studies showed myocilin promotes cell migration and phagocytic activities of human trabecular meshwork cells[40,59,60]. In our work, overexpressed MYOC in TDMs elevated phagocytic activities and migration, which induced more intracellular cryptococci and enhanced effects of “Trojan Horse”. These indicated MYOC may the effector for cryptococcoses secondary to HIV/AIDS.

In conclusion, our findings therefore described global miRNA-mRNA regulatory responses during *C. neoformans* in primate and rodent animal models, which serves as a clinically relevant database for fundamental and clinical research. We highlight the importance of MYOC and cytoskeleton pathways during *Cryptococcus* meningoencephalitis and underscores their critical functions in the formation of “Trojan Horse” (Figure 7). This study provides critical roles of cytoskeleton on fungal CNS invasion, reveals the directly reasons for the high prevalence of cryptococcal meningitis in HIV/AIDS, and facilitates possibilities for novel anti-fungal drugs development.

**Figure 7.**
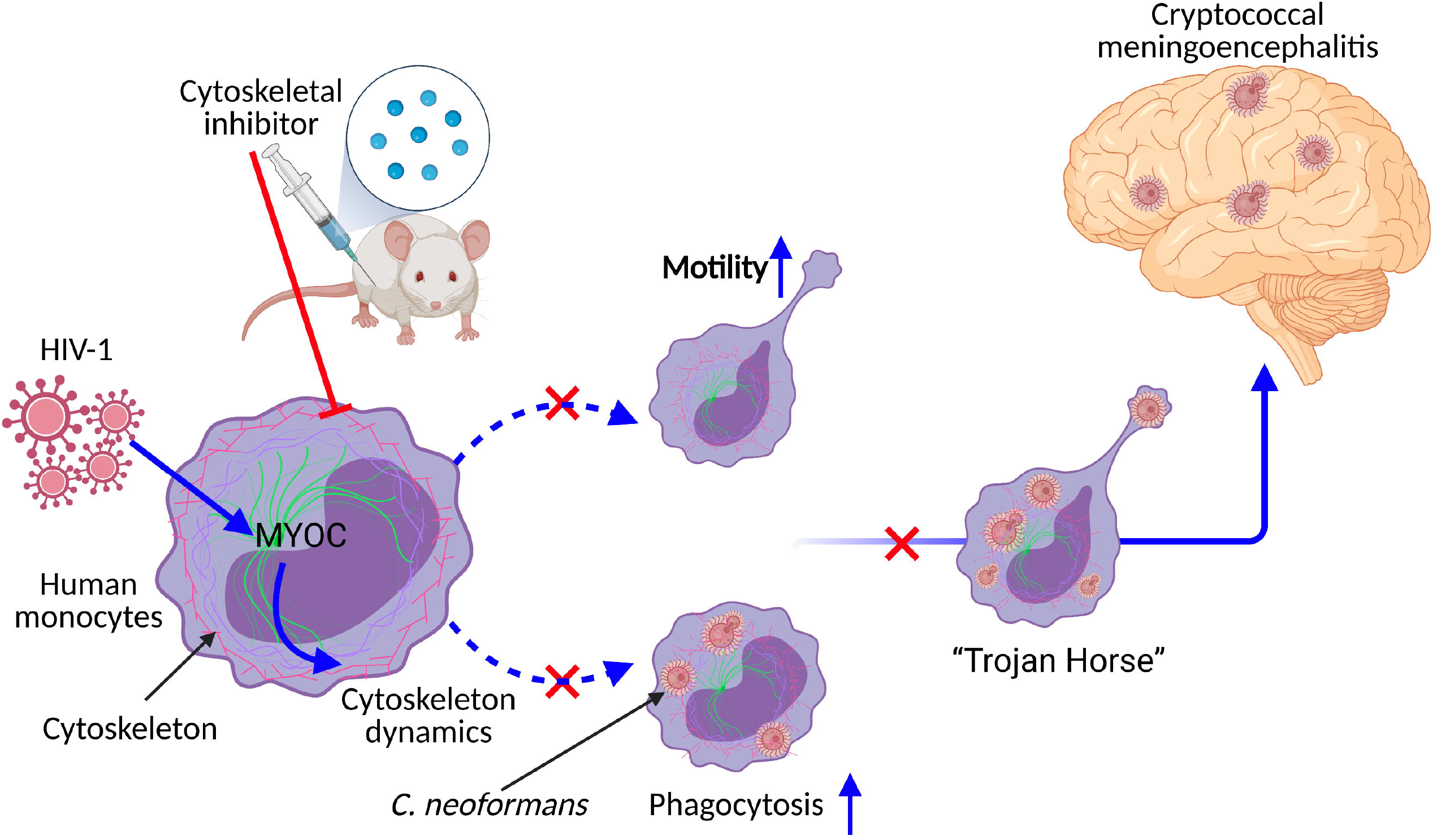
Conceptual graph of HIV triggered cryptococcal meningoencephalitis. HIV-1 infections trigger cytoskeleton dynamics by induction of MYOC protein, enhance macrophage “Trojan Horse” and induce cryptococcus meningoencephalitis (shown in blue lines). Inhibitions of cytoskeleton pathways or MYOC dampen “Trojan Horse” and decrease the number of cryptococci in the brain (shown in red lines). Figure 7 was created with BioRender.com

## Acknowledgments

The authors would like to thank Pro. Tongbao Liu from Southwest University, China, for kindly providing GFP-labelled *C. neoformans* strain. This research was supported by the CAMS Innovation Fund for Medical Sciences (2019-I2M-5-027 to HS and XH), the China Postdoctoral Science Foundation (2021M693520 to HL), and the National Natural Science Foundation of China (31870140 to CD and 81801989 to TS).

## Disclosure statement

No potential conflict of interest was reported by the authors.

## Supporting information

**Figure S1.**
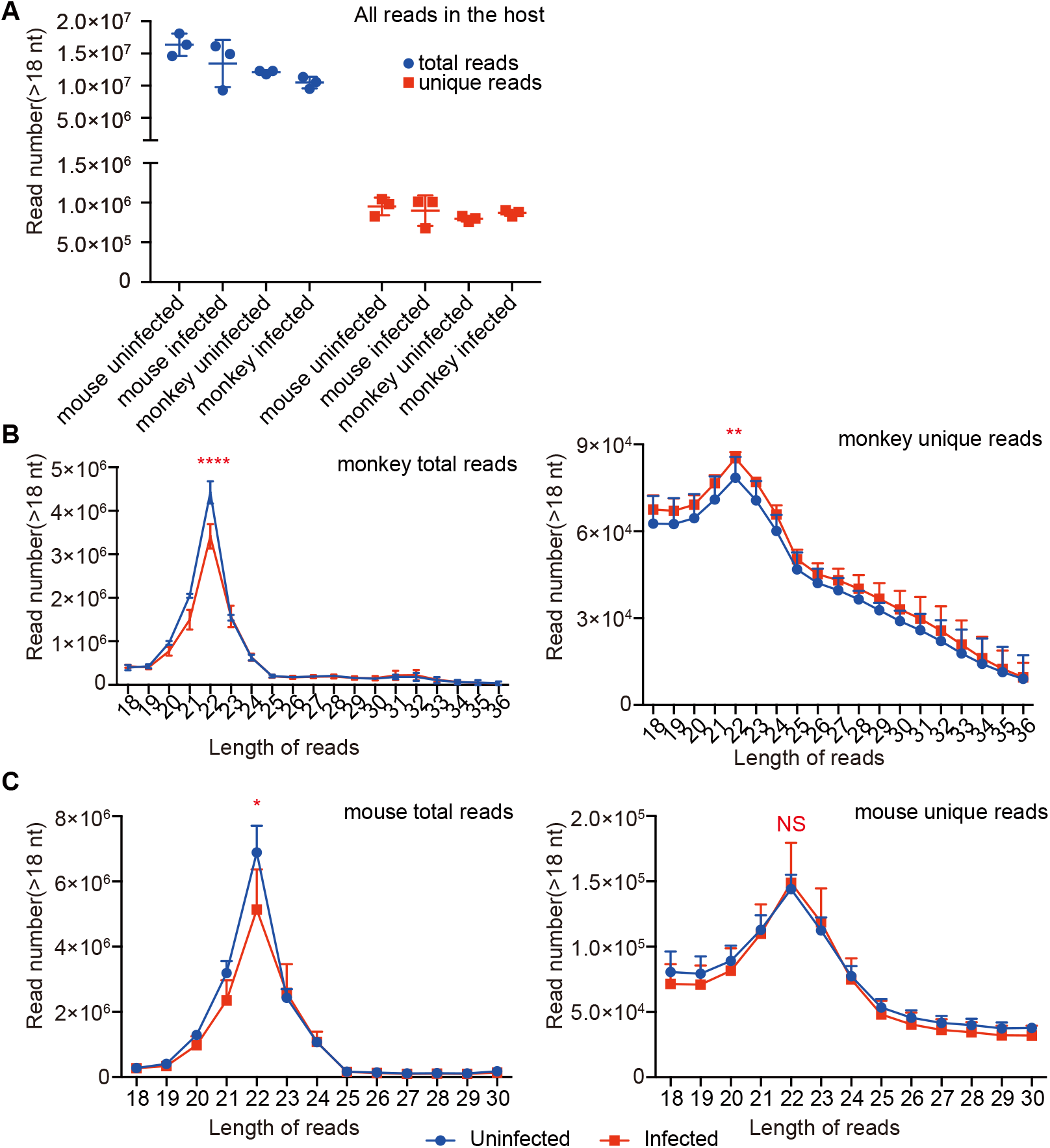
Read number of miRNA-Seq. **A**. Total and unique read number of miRNA-Seq. **B**. Distribution of different lengths of nucleotides from macaques. **C**. Distribution of different lengths of nucleotides from mice.

**Figure S2.**
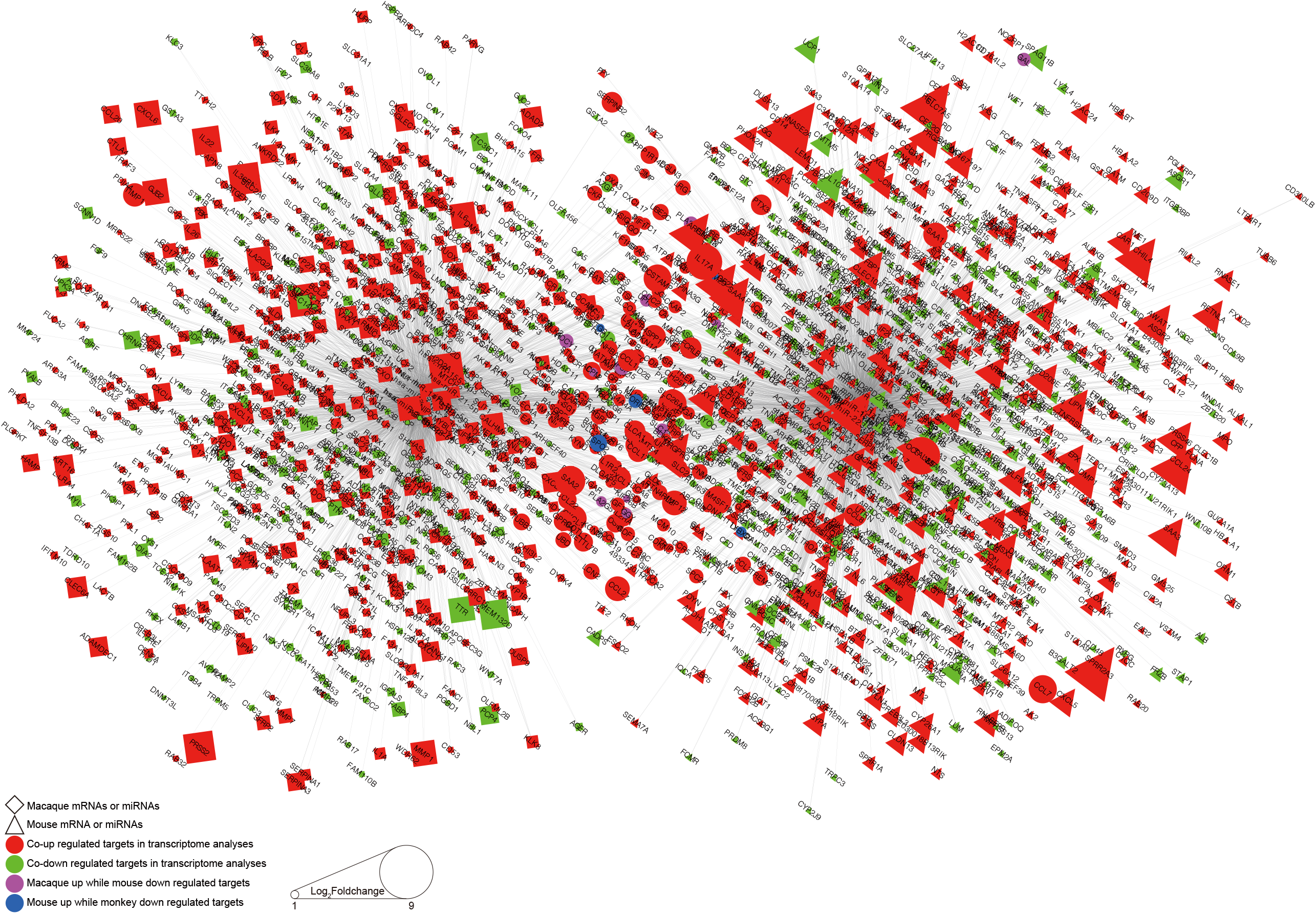
Network of miRNA-mRNA during cryptococcal pneumonia.

**Figure S3.**
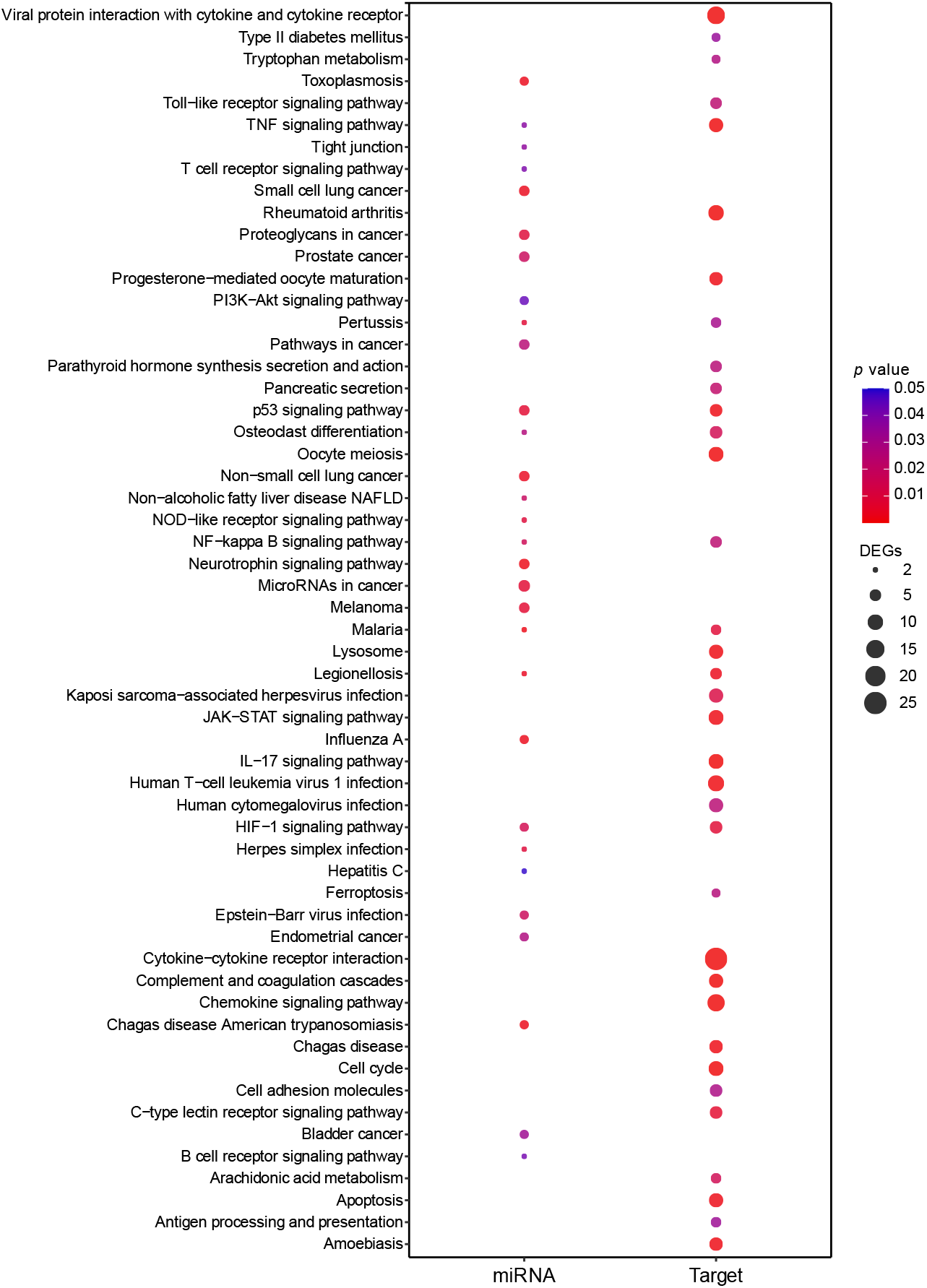
KEGG analysis of co-regulated miRNAs and targeted mRNAs.

**Figure S4.**
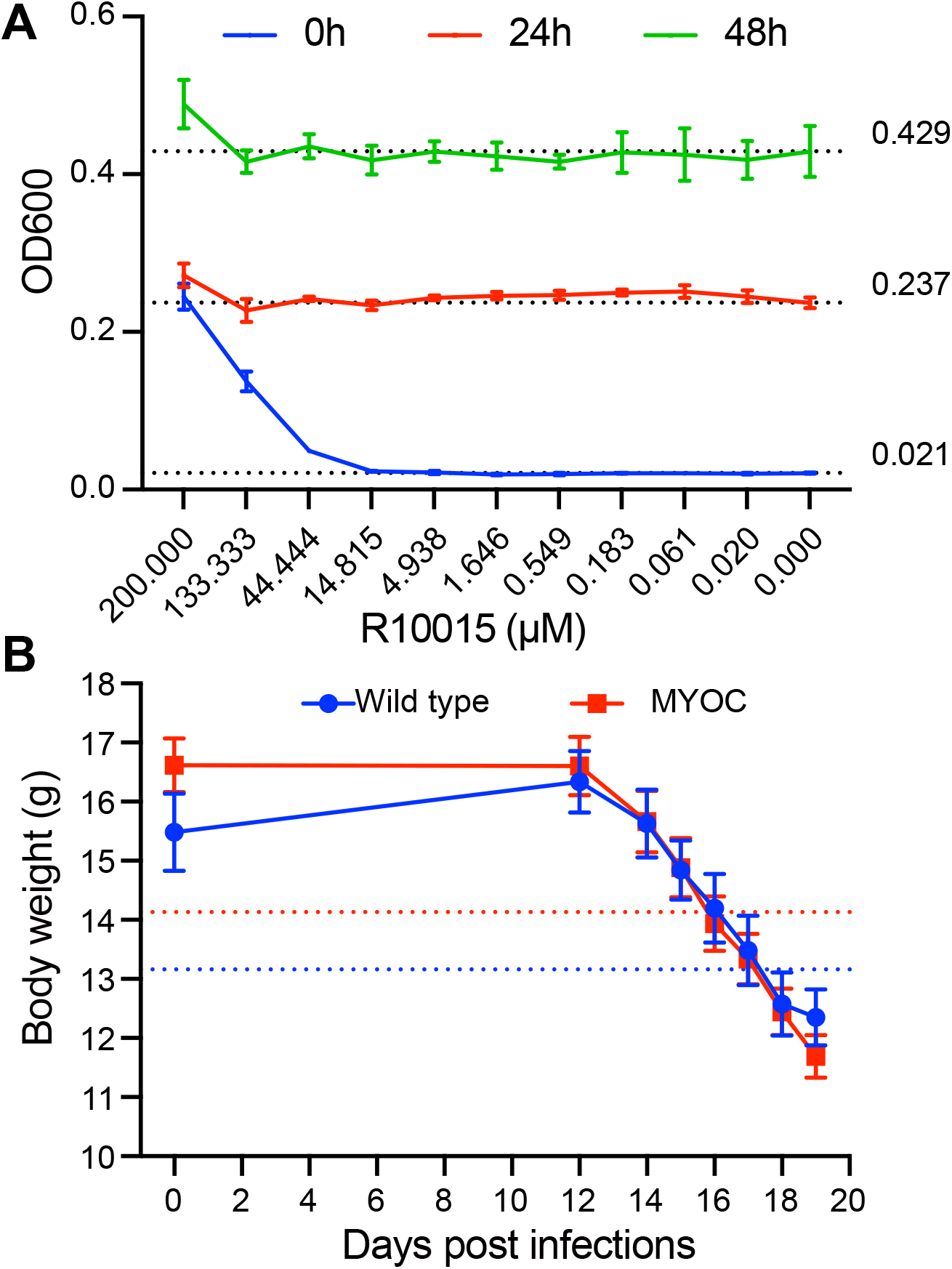
Growth curve of *C. neoformans* and body weight loss of mice in survival rates. **A. Growth curve of H99 in different concentrations of R10015**. **B. Body weight of mice**. Blue line indicated wild type control group and red line MYOC transgenic mice. Dash dot lines represent 85% of the initial average body weight.

**Table S1**. Differentially expressed miRNAs in macaque and mouse lung tissues during *C. neoformans* infections.

**Table S2**. GO and KEGG analyses of 8 co-regulated miRNAs in response to *C. neoformans* infections.

**Table S3**. GO and KEGG analyses of target genes for co-regulated 8 miRNAs in mouse and macaque.

**Table S4**. Primers used in the study.

## Notes

### Competing Interest Statement

The authors have declared no competing interest.

## Reference

1. Iyer KR, Revie NM, Fu C, et al. Treatment strategies for cryptococcal infection: challenges, advances and future outlook. Nat Rev Microbiol. 2021 Jul;19(7):454–466.

2. Stott KE, Loyse A, Jarvis JN, et al. Cryptococcal meningoencephalitis: time for action. Lancet Infect Dis. 2021 Sep;21(9):e259–e271.

3. Mpoza E, Rajasingham R, Tugume L, et al. Cryptococcal Antigenemia in Human Immunodeficiency Virus Antiretroviral Therapy-Experienced Ugandans With Virologic Failure. Clin Infect Dis. 2020 Oct 23;71(7):1726–1731.

4. Chen M, Xu N, Xu J. Cryptococcus Neoformans Meningitis Cases Among China’s HIV-Infected Population may have been Severely Under-Reported. Mycopathologia. 2020 Dec;185(6):971–974.

5. MacDougall L, Kidd SE, Galanis E, et al. Spread of Cryptococcus gattii in British Columbia, Canada, and detection in the Pacific Northwest, USA. Emerg Infect Dis. 2007 Jan;13(1):42–50.

6. Huang C, Tsui CKM, Chen M, et al. Emerging Cryptococcus gattii species complex infections in Guangxi, southern China. PLoS Negl Trop Dis. 2020 Aug;14(8):e0008493.

7. Bielska E, Sisquella MA, Aldeieg M, et al. Pathogen-derived extracellular vesicles mediate virulence in the fatal human pathogen Cryptococcus gattii. Nat Commun. 2018 Apr 19;9(1):1556.

8. Fang W, Zhang L, Liu J, et al. Tuberculosis/cryptococcosis co-infection in China between 1965 and 2016. Emerg Microbes Infect. 2017 Aug 23;6(8):e73.

9. Shen Y, Zheng F, Sun D, et al. Epidemiology and clinical course of COVID-19 in Shanghai, China. Emerg Microbes Infect. 2020 Dec;9(1):1537–1545.

10. Cafardi J, Haas D, Lamarre T, et al. Opportunistic Fungal Infection Associated With COVID-19. Open Forum Infect Dis. 2021 Jul;8(7):ofab016.

11. Song G, Liang G, Liu W. Fungal Co-infections Associated with Global COVID-19 Pandemic: A Clinical and Diagnostic Perspective from China. Mycopathologia. 2020 Aug;185(4):599–606.

12. Traver EC, Sánchez MM. Pulmonary aspergillosis and cryptococcosis as a complication of COVID-19. Medical Mycology Case Reports. 2022.

13. Charlier C, Nielsen K, Daou S, et al. Evidence of a role for monocytes in dissemination and brain invasion by Cryptococcus neoformans. Infect Immun. 2009 Jan;77(1):120–7.

14. Santiago-Tirado FH, Onken MD, Cooper JA, et al. Trojan Horse Transit Contributes to Blood-Brain Barrier Crossing of a Eukaryotic Pathogen. mBio. 2017 Jan 31;8(1).

15. Scherer AK, Blair BA, Park J, et al. Redundant Trojan horse and endothelial-circulatory mechanisms for host-mediated spread of Candida albicans yeast. PLoS Pathog. 2020 Aug;16(8):e1008414.

16. Sorrell TC, Juillard PG, Djordjevic JT, et al. Cryptococcal transmigration across a model brain blood-barrier: evidence of the Trojan horse mechanism and differences between Cryptococcus neoformans var. grubii strain H99 and Cryptococcus gattii strain R265. Microbes Infect. 2016 Jan;18(1):57–67.

17. Strickland AB, Shi M. Mechanisms of fungal dissemination. Cell Mol Life Sci. 2021 Jan 15.

18. Alves Soares E, Lazera MDS, Wanke B, et al. Mortality by cryptococcosis in Brazil from 2000 to 2012: A descriptive epidemiological study. PLoS Negl Trop Dis. 2019 Jul;13(7):e0007569.

19. Akaihe CL, Nweze EI. Epidemiology of Cryptococcus and cryptococcosis in Western Africa. Mycoses. 2021 Jan;64(1):4–17.

20. Rajasingham R, Smith RM, Park BJ, et al. Global burden of disease of HIV-associated cryptococcal meningitis: an updated analysis. Lancet Infect Dis. 2017 Aug;17(8):873–881.

21. Walsh D, Naghavi MH. Exploitation of Cytoskeletal Networks during Early Viral Infection. Trends Microbiol. 2019 Jan;27(1):39–50.

22. Li H, Li Y, Sun T, et al. Unveil the transcriptional landscape at the Cryptococcus-host axis in mice and nonhuman primates. PLoS Negl Trop Dis. 2019 Jul;13(7):e0007566.

23. Chen SHM, Stins MF, Huang SH, et al. Cryptococcus neoformans induces alterations in the cytoskeleton of human brain microvascular endothelial cells. J Med Microbiol. 2003 Nov;52(Pt 11):961-970.

24. Li H, Li Y, Sun T, et al. Integrative Proteome and Acetylome Analyses of Murine Responses to Cryptococcus neoformans Infection. Front Microbiol. 2020;11:575.

25. Li Y, Li H, Sui M, et al. Fungal acetylome comparative analysis identifies an essential role of acetylation in human fungal pathogen virulence. Commun Biol. 2019;2:154.

26. Holmer SM, Evans KS, Asfaw YG, et al. Impact of surfactant protein D, interleukin-5, and eosinophilia on Cryptococcosis. Infect Immun. 2014 Feb;82(2):683–93.

27. Muller U, Stenzel W, Kohler G, et al. IL-13 induces disease-promoting type 2 cytokines, alternatively activated macrophages and allergic inflammation during pulmonary infection of mice with Cryptococcus neoformans. J Immunol. 2007 Oct 15;179(8):5367–77.

28. Angkasekwinai P, Sringkarin N, Supasorn O, et al. Cryptococcus gattii infection dampens Th1 and Th17 responses by attenuating dendritic cell function and pulmonary chemokine expression in the immunocompetent hosts. Infect Immun. 2014 Sep;82(9):3880–90.

29. Marais S, Meintjes G, Lesosky M, et al. Interleukin-17 mediated differences in the pathogenesis of HIV-1-associated tuberculous and cryptococcal meningitis. AIDS. 2016 Jan 28;30(3):395–404.

30. Murdock BJ, Huffnagle GB, Olszewski MA, et al. Interleukin-17A enhances host defense against cryptococcal lung infection through effects mediated by leukocyte recruitment, activation, and gamma interferon production. Infect Immun. 2014 Mar;82(3):937–48.

31. Wozniak KL, Hardison SE, Kolls JK, et al. Role of IL-17A on resolution of pulmonary C. neoformans infection. PLoS One. 2011 Feb 17;6(2):e17204.

32. Chen H, Jin Y, Chen H, et al. MicroRNA-mediated inflammatory responses induced by Cryptococcus neoformans are dependent on the NF-kappaB pathway in human monocytes. Int J Mol Med. 2017 Jun;39(6):1525–1532.

33. Liu M, Zhang Z, Ding C, et al. Transcriptomic Analysis of Extracellular RNA Governed by the Endocytic Adaptor Protein Cin1 of Cryptococcus deneoformans. Front Cell Infect Microbiol. 2020;10:256.

34. Zhang L, Zhang K, Fang W, et al. CircRNA-1806 Decreases T Cell Apoptosis and Prolongs Survival of Mice After Cryptococcal Infection by Sponging miRNA-126. Front Microbiol. 2020;11:596440.

35. Jin Y, Yao G, Wang Y, et al. MiR-30c-5p mediates inflammatory responses and promotes microglia survival by targeting eIF2alpha during Cryptococcus neoformans infection. Microb Pathog. 2020 Apr;141:103959.

36. Busk PK. A tool for design of primers for microRNA-specific quantitative RT-qPCR. BMC Bioinformatics. 2014 Jan 28;15:29.

37. Sticht C, De La Torre C, Parveen A, et al. miRWalk: An online resource for prediction of microRNA binding sites. PLoS One. 2018;13(10):e0206239.

38. Kern F, Fehlmann T, Solomon J, et al. miEAA 2.0: integrating multi-species microRNA enrichment analysis and workflow management systems. Nucleic Acids Res. 2020 Jul 2;48(W1):W521-W528.

39. Yi F, Guo J, Dabbagh D, et al. Discovery of Novel Small-Molecule Inhibitors of LIM Domain Kinase for Inhibiting HIV-1. J Virol. 2017 Jul 1;91(13).

40. Kwon HS, Tomarev SI. Myocilin, a Glaucoma-Associated Protein, Promotes Cell Migration Through Activation of Integrin-Focal Adhesion Kinase-Serine/Threonine Kinase Signaling Pathway. Journal of Cellular Physiology. 2011 Dec;226(12):3392–3402.

41. Kim JC, Crary B, Chang YC, et al. Cryptococcus neoformans activates RhoGTPase proteins followed by protein kinase C, focal adhesion kinase, and ezrin to promote traversal across the blood-brain barrier. J Biol Chem. 2012 Oct 19;287(43):36147–57.

42. Zhang C, Chen F, Liu X, et al. Gliotoxin Induces Cofilin Phosphorylation to Promote Actin Cytoskeleton Dynamics and Internalization of Aspergillus fumigatus Into Type II Human Pneumocyte Cells. Front Microbiol. 2019;10:1345.

43. Kanjanapruthipong T, Sukphopetch P, Reamtong O, et al. Cytoskeletal Alteration Is an Early Cellular Response in Pulmonary Epithelium Infected with Aspergillus fumigatus Rather than Scedosporium apiospermum. Microbial ecology. 2021:1–20.

44. Johnston SA, May RC. The human fungal pathogen Cryptococcus neoformans escapes macrophages by a phagosome emptying mechanism that is inhibited by Arp2/3 complex-mediated actin polymerisation. PLoS Pathog. 2010 Aug 12;6(8):e1001041.

45. Tang DD, Gerlach BD. The roles and regulation of the actin cytoskeleton, intermediate filaments and microtubules in smooth muscle cell migration. Respir Res. 2017 Apr 8;18(1):54.

46. Meyerovitch J, Farfel Z, Sack J, et al. Oral administration of vanadate normalizes blood glucose levels in streptozotocin-treated rats. Characterization and mode of action. J Biol Chem. 1987 May 15;262(14):6658–62.

47. Kogan TV, Jadoun J, Mittelman L, et al. Involvement of secreted Aspergillus fumigatus proteases in disruption of the actin fiber cytoskeleton and loss of focal adhesion sites in infected A549 lung pneumocytes. J Infect Dis. 2004 Jun 1;189(11):1965–73.

48. He S, Fu Y, Guo J, et al. Cofilin hyperactivation in HIV infection and targeting the cofilin pathway using an anti-alpha4beta7 integrin antibody. Sci Adv. 2019 Jan;5(1):eaat7911.

49. Montoya MC, Magwene PM, Perfect JR. Associations between Cryptococcus Genotypes, Phenotypes, and Clinical Parameters of Human Disease: A Review. J Fungi (Basel). 2021 Mar 30;7(4).

50. Molloy SF, Kanyama C, Heyderman RS, et al. Antifungal Combinations for Treatment of Cryptococcal Meningitis in Africa. N Engl J Med. 2018 Mar 15;378(11):1004–1017.

51. Meya D, Rajasingham R, Nalintya E, et al. Preventing Cryptococcosis-Shifting the Paradigm in the Era of Highly Active Antiretroviral Therapy. Curr Trop Med Rep. 2015;2(2):81–89.

52. Tugume L, Rhein J, Hullsiek KH, et al. HIV-associated cryptococcal meningitis occurring at relatively higher CD4 counts. The Journal of infectious diseases. 2019;219(6):877–883.

53. Fisher JF, Valencia-Rey PA, Davis WB. Pulmonary Cryptococcosis in the Immunocompetent Patient-Many Questions, Some Answers. Open Forum Infect Dis. 2016 Sep;3(3):ofw167.

54. Yoder A, Yu D, Dong L, et al. HIV envelope-CXCR4 signaling activates cofilin to overcome cortical actin restriction in resting CD4 T cells. Cell. 2008 Sep 5;134(5):782–92.

55. Seoane PI, Taylor-Smith LM, Stirling D, et al. Viral infection triggers interferon-induced expulsion of live Cryptococcus neoformans by macrophages. PLoS Pathog. 2020 Feb;16(2):e1008240.

56. Mertts M, Garfield S, Tanemoto K, et al. Identification of the region in the N-terminal domain responsible for the cytoplasmic localization of Myoc/Tigr and its association with microtubules. Lab Invest. 1999 Oct;79(10):1237–45.

57. Sohn S, Hur W, Joe MK, et al. Expression of wild-type and truncated myocilins in trabecular meshwork cells: their subcellular localizations and cytotoxicities. Investigative ophthalmology & visual science. 2002;43(12):3680–3685.

58. Shen X, Koga T, Park BC, et al. Rho GTPase and cAMP/protein kinase A signaling mediates myocilin-induced alterations in cultured human trabecular meshwork cells. J Biol Chem. 2008 Jan 4;283(1):603–612.

59. Wentz-Hunter K, Kubota R, Shen X, et al. Extracellular myocilin affects activity of human trabecular meshwork cells. J Cell Physiol. 2004 Jul;200(1):45–52.

60. Wentz-Hunter K, Shen X, Okazaki K, et al. Overexpression of myocilin in cultured human trabecular meshwork cells. Exp Cell Res. 2004 Jul 1;297(1):39–48.

